# The macronutrient content of a meal modulates subsequent ‘dessert’ intake, via Fibroblast Growth Factor 21 (FGF21)

**DOI:** 10.1101/2023.07.07.547884

**Authors:** Chih-Ting Wu, Karlton R. Larson, Landon C. Sims, Karen K. Ryan

**Affiliations:** Department of Neurobiology, Physiology and Behavior, College of Biological Sciences, University of California, Davis, CA, 95616, USA

**Keywords:** fibroblast growth factor 21, macronutrient, preload, snack, dessert, hedonic, protein, sucrose

## Abstract

Pharmacological administration of Fibroblast growth factor 21 (FGF21) alters food choice, including that it decreases the consumption of sucrose and other sweet tastants. Conversely, endogenous secretion of FGF21 by the liver is modulated by diet, such that plasma FGF21 is increased after eating foods that have a low dietary protein: total energy (P: E) ratio. Together, these findings suggest a strategy to promote healthy eating, in which the macronutrient content of a pre-load meal could reduce the later consumption of sweet desserts. Here, we tested the prediction that individuals eating a low P: E pre-load meal, and next offered a highly palatable sweet ‘dessert’, would eat less of the sugary snack compared to controls, due to increased FGF21 signaling. In addition to decreasing sweet intake, FGF21 increases the consumption of dietary protein. Thus, we predicted that individuals eating a low protein pre-load meal, and subsequently offered a very high-protein pellet as ‘dessert’ or snack, would eat more of the high protein pellet compared to controls, and that this depends on FGF21. We tested this in C57Bl/6J, and liver-specific FGF21-null (FGF21^ΔL^) null male and female mice and littermate controls. Contrary to expectation, eating a low protein pre-load did not reduce the later consumption of a sweet solution in either males or females, despite robustly increasing plasma FGF21. Rather, eating the low protein pre-load increased later consumption of a high protein pellet. This was more apparent among males and was abrogated in the FGF21^ΔL^ mice. We conclude that physiologic induction of hepatic FGF21 by a low protein pre-load is not sufficient to reduce later consumption of sweet dessert, though it effectively increases the subsequent intake of dietary protein in male mice.

## Introduction

It is generally considered that greater consumption of sugar sweetened beverages and sweet desserts like soda, cakes, and cookies, is associated with overweight and obesity [1,2], and increases the risk of non-alcoholic fatty liver disease [3], diabetes [4,5], and other cardiometabolic diseases [4–7]. Dietary recommendations for weight management and the prevention of chronic diseases have long included the avoidance of added sugars, highly-processed sweet treats, and sugar sweetened beverages, but compliance with these recommendations is generally poor [8,9] and overall consumption of dietary sugars continues to rise [10]. Nutritional and/or behavioral strategies that promote healthy eating, by reducing the motivation to consume sugary treats and increasing compliance with dietary recommendations, could have a broad impact on public health.

Our lab and others recently demonstrated that the hormone fibroblast growth factor-21 (FGF21) is robustly induced in response to dietary protein dilution (low P: E diet) in rodents [11–16] and humans [13,15], and acts in the brain to control feeding behavior [11,17–20]. Specifically, endogenous secretion of FGF21 from the liver is elevated when animals consume simple sugars [19,21,22], or when they otherwise eat a diet that is low in dietary protein relative to total energy (low P: E) [11–15,23]. In turn, pharmacological administration of FGF21 decreases the consumption of sucrose and other sweet tastants and it increases the consumption of dietary protein [17–19,24]. Taken together, this suggests a mechanism by which the macronutrient content of a pre-load meal may influence the subsequent consumption of a dessert or snack, via FGF21. Specifically, we hypothesized that individuals consuming a pre-load meal having a low P:E ratio will reduce the subsequent intake of a less healthy sweet dessert, compared to controls. Conversely, and consistent with the prior literature [11,25,26], we expect these individuals will increase the consumption of an arguably healthier protein-rich ‘dessert’, despite that very high-protein foods are not typically considered highly palatable or rewarding. Lastly, we hypothesize that these changes in feeding behavior require FGF21 signaling.

## Materials and Methods

### Animals

Age-matched male and female C57Bl6J mice, liver-specific FGF21-null (FGF21^ΔL^) mice, and control littermates, 9 to 15 weeks of age, were used in this study. C57BI/6J mice were obtained from The Jackson Laboratory or were bred in-house up to 2 generations removed from the founders. FGF21^ΔL^ mice and littermate controls (FGF21^Δfl/fl^) were generated by breeding transgenic mice expressing Cre-recombinase under Albumin promoter (#003574, Jackson Labs) with FGF21-floxed mice (#022361, Jackson Labs) and maintained on a C57Bl/6J background in our lab. The founders of our colony were a generous gift from Dr. Brian DeBosch, Washington University in St. Louis. Mice were singly housed on 12hr light/dark cycle in a temperature (20ºC to 22ºC) and humidity-controlled vivarium with *ad libitum* access to food and water unless otherwise noted. In all experiments, mice were counterbalanced to dietary treatment groups by body weight. All animal experiments were approved by the Institutional Care and Use Committees of the University of California, Davis.

### Diets

Mice were maintained on standard chow diet (Harlan, catalog #5008, Madison, WI) until the study began. Sucrose (Sigma, St. Louis, MO) was dissolved in distilled water to make a sucrose solution with the indicated concentration. Solid sucrose pellets (F0023, 45 mg unflavored) were manufactured by Bio-Serv (Frenchtown, NJ). Purified, pelleted normal (18%), high (36%), and low (4%) protein diets (D11092301, D11051801, and D11092304) were manufactured by Research Diets (New Brunswick, NJ). Full nutritional details can be found in Table 1.

**Table 1:**
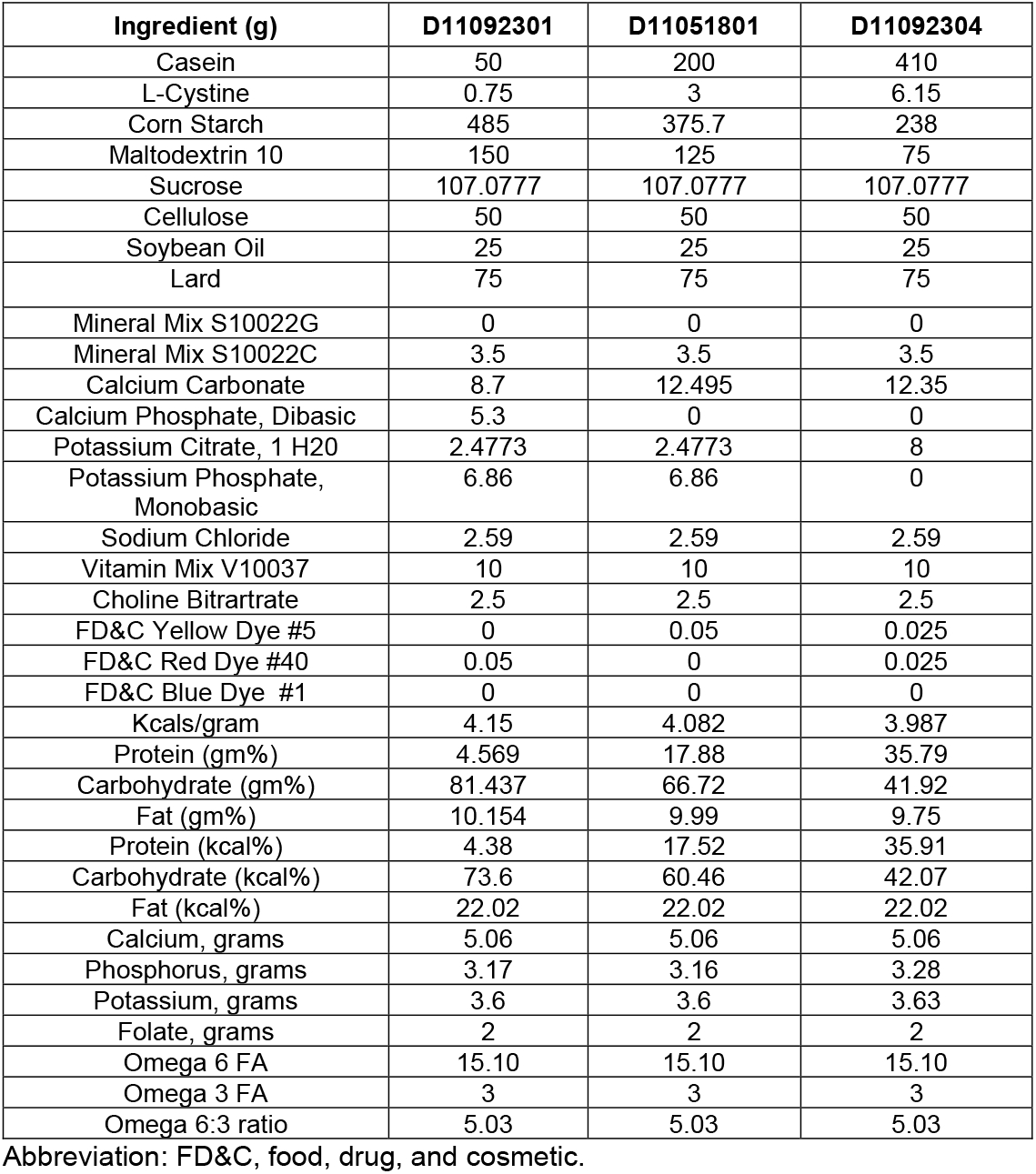
Diet composition for pelleted diets

### Dessert feeding behavior

We used a previously-established “dessert effect” paradigm (e.g., [27–31]) to examine the consumption of palatable foods by calorically-sated mice (Fig. 1). This protocol mimics the common experience that, even after eating an initial meal or pre-load to caloric satiation, individuals may be motivated to further consume a palatable dessert.

**Figure 1.**
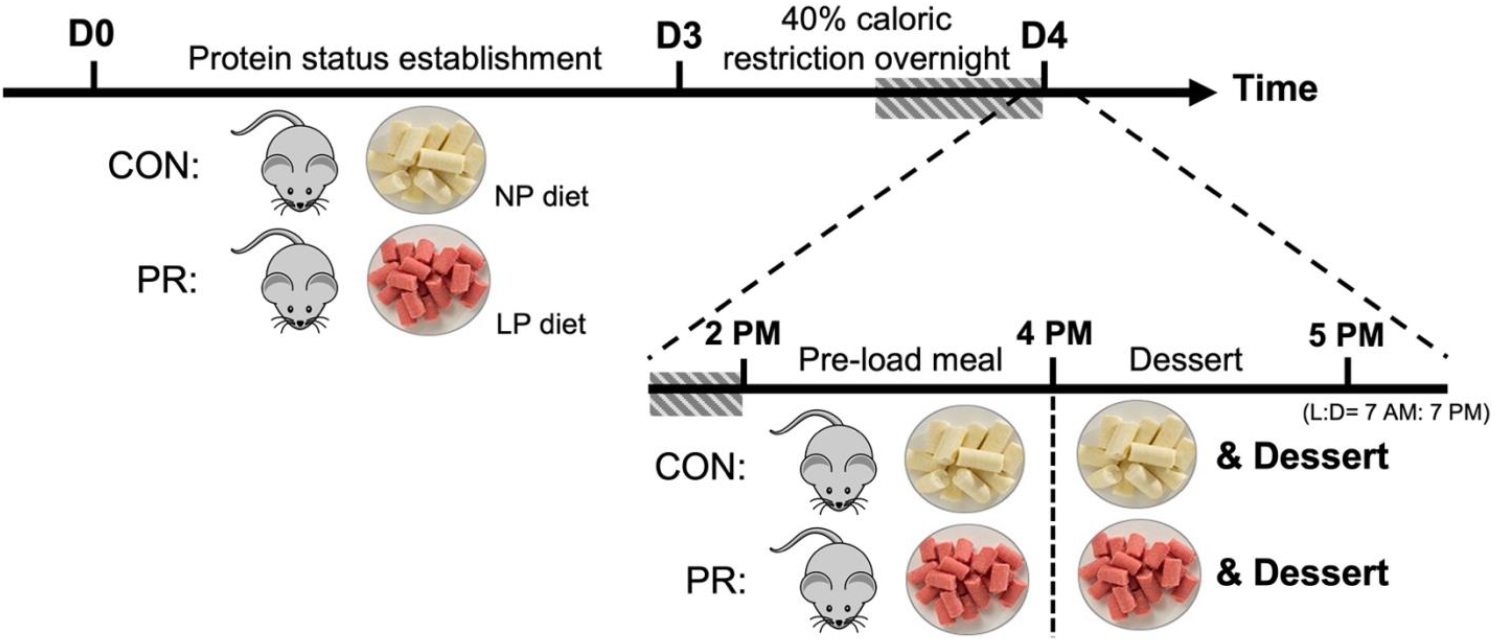
Experimental design of the “Dessert Effect” test. Male and female mice were maintained on NP and LP diets for 3 days to establish control (CON) and protein restricted (PR) nutritional status, respectively. On the night before behavioral testing, mice were energy-restricted to 60% of their daily caloric intake, to motivate and synchronize food intake during the test. On test day, mice were fed an initial 2-hour preload. This consisted of *ad libitum* access to the previously assigned LP or NP maintenance diet. Next, we additionally provided *ad libitum* access to a ‘dessert’ diet for 1-hour.

Mice were maintained on one of two purified, controlled diets that differ in macronutrient content but are matched for sweetness and caloric density. The low-protein diet (LP) contained 4% PRO, 74% CHO, 22% fat by kcal, whereas the control normal-protein diet (NP) contained 18% PRO, 60% CHO, 22% fat by kcal (Table 1). Mice were maintained on these diets for 3 days to establish protein restricted (PR) and control (CON) status, respectively. On the night before behavioral testing, mice were energy-restricted to 60% of their daily caloric intake, to motivate meal feeding and synchronize food intake during the “dessert effect” test. On test day, mice were fed an initial 2-hour ‘pre-load’ beginning at ZT 7. This consisted of *ad libitum* access to the previously assigned LP or NP maintenance diet. Next, beginning at ZT 9, we additionally provided *ad libitum* access to a ‘dessert’ diet for 1-hour. This dessert consisted of either sucrose solution (Fig. 2) or a high-protein pelleted diet (36% PRO, 42% CHO, 22% fat) (Figs. 3-4). Note: mice were briefly exposed to the ‘dessert’ diets approximately 1 week before behavioral testing, to prevent neophobia.

**Figure 2.**
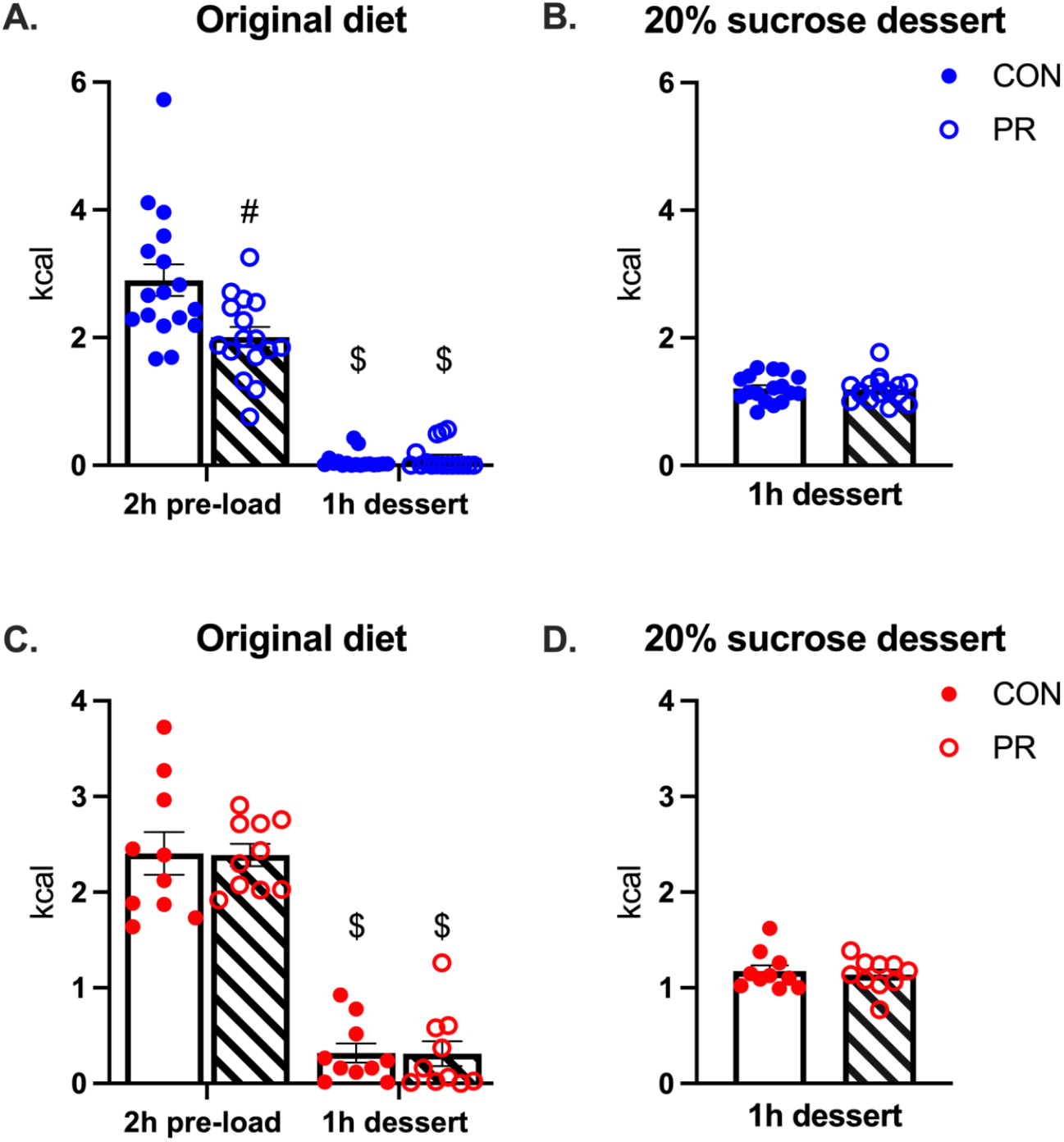
Protein status of a pre-load meal did not affect subsequent consumption of 20% sucrose dessert. Male (A-B) and female (C-D) CON and PR mice consumed the majority of their caloric intake from original maintenance diets during the 2h pre-load meal and ate relatively little pelleted diet during the ‘dessert’ phase of the experiment [$: p<0.0001 versus 2h pre-load; #: p<0.001 versus CON] (A, C). There was no effect of pre-load diet on the consumption of 20% sucrose dessert (B, D). Analyses made by two-way ANOVA with Tukey *post hoc* tests (A,C) or by T-test (B,D). Data are shown as mean ± SEM, n= 9-17 mice per group.

### Plasma analysis

In a separate cohort of C57/BL6J mice, we confirmed that 3 days of NP vs LP-feeding induces FGF21 secretion. We collected blood from the tip of the tail, into chilled EDTA-coated tubes and centrifuged at 3000 g for 20 minutes at 4°C. Plasma was stored at -80°C for later use. FGF21 was measured by using a commercially available enzyme-linked immunosorbent assay kit (rat/mouse FGF21 enzyme-linked immunosorbent assay kit; Millipore, Billerica, MA), according to the manufacturer’s instructions.

### Statistical analysis

Data were analyzed using GraphPad Prism by the appropriate model analysis of variance (ANOVA) or t-test, as indicated. Multiple comparisons were made using Tukey *post hoc* tests. In all cases, α= 0.05. Graphs were created using GraphPad Prism. Data are presented as means +/-SEM unless otherwise noted.

## Results

### Protein content of the pre-load did not reduce the consumption of sweet dessert in males and females

For both males (Fig. 2A-B) and females (Fig. 2C-D), we found both protein-restricted and control mice consumed most of their total caloric intake from maintenance diet during the 2-h preload and ate relatively little pelleted diet during the ‘dessert’ phase of the experiment [p (time)< 0.001, Tukey’s posthoc, p< 0.001 (Fig. 2A, C)]. Contrary to expectation, however, there was no effect of pre-load diet on the subsequent consumption of sucrose dessert [T-test, (Fig. 2B, D)]. We repeated this study in two additional cohorts, using more (30%) and less (10%) concentrated sucrose solutions to account for variance in palatability. Similarly, no difference in sucrose consumption was observed in either males or females (Supplementary Fig. 1&2). Thus, a LP pre-load did not suppress the consumption of the subsequent sucrose dessert in mice.

Pharmacologic delivery of FGF21 inhibits the consumption of sweet solutions, including 10% sucrose and 0.2% sucralose [18,19,32]. FGF21 also increases water intake [33,34], presenting a potential confound in our experimental design, because the mice had both sucrose and water available during the test. Therefore, we repeated the experiment for a fourth time, using solid sucrose pellets instead of dissolving the sucrose in water. Still, there was no effect of protein status on the consumption of sucrose pellets as ‘dessert’ (Supplementary Fig. 3). Taken together, these data do not support the initial hypothesis for this study—that protein content of a pre-load meal will reduce the subsequent consumption of a sweet dessert.

### Protein-restricted males, but not females, significantly increased the consumption of a high-protein ‘dessert’

As before, for both male (Fig. 3A-B) and female (Fig. 3C-D) mice, we found both protein-restricted mice and controls consumed most of their total caloric intake from maintenance diet, during the 2-h preload meal [p (time)< 0.001, Tukey’s posthoc, p< 0.001 (Fig. 3A, C)]. Consistent with our prediction, we found protein restricted male mice ate significantly more of the high-protein ‘dessert’ or snack compared to controls [T-test, p< 0.001 (Fig. 3B)]. However, we did not observe an effect of protein restriction on dessert intake in females [T-test, p=0.3296 (Fig. 3D)]. To determine a potential mechanism for this differential response, we measured plasma FGF21 in CON and PR male and female littermates. Notably, although protein restriction significantly enhanced plasma FGF21 in both sexes, the magnitude of this increase depended on sex [2-way ANOVA; p (diet x sex) < 0.05]. PR male mice had significantly higher plasma FGF21 than PR females (Fig. 3E).

**Figure 3.**
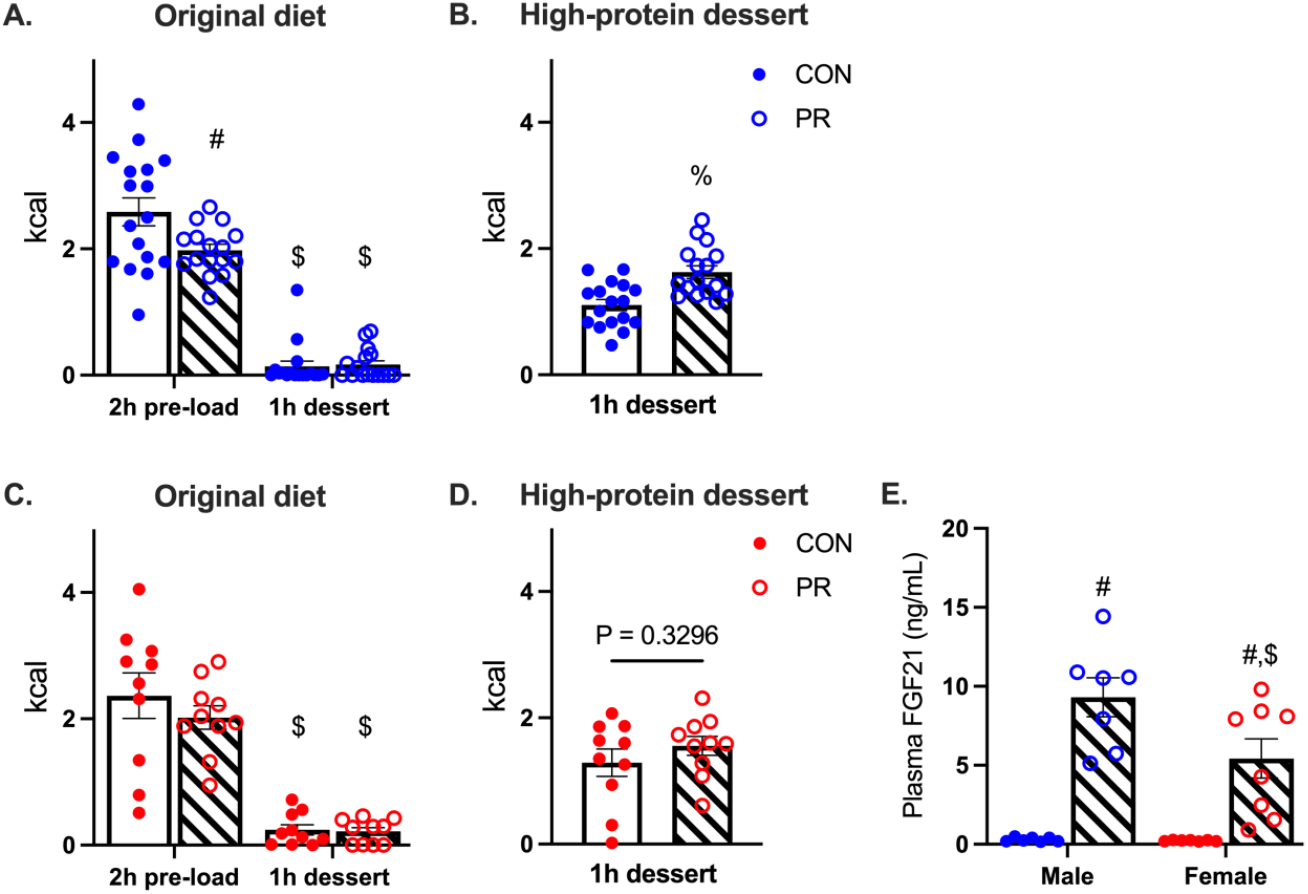
Protein content of a pre-load meal increased subsequent consumption of a high protein ‘dessert’. Both male (A-B) and female (C-D) CON and PR mice consumed most of their caloric intake from original maintenance diets during the 2h pre-load meal and ate little of the maintenance diet during the ‘dessert’ phase of the test [$: p<0.0001 versus 2h pre-load; #: p<0.01 versus CON] (A, C). PR male mice ate significantly more high-protein dessert compared to CON mice [T-test, %: p<0.001] (B), but this was not observed in PR females (D). Plasma FGF21 was markedly enhanced after 3-days of dietary protein restriction in both males and females, but the magnitude of this effect depended on sex. PR males had significantly higher plasma FGF21 level compared to PR female littermates [#: p<0.001 versus CON; $: p<0.05 versus male PR] (E). Analyses made by two-way ANOVA with Tukey *post hoc* tests (A, n = 16-17 mice/ group; C, n = 10 mice/ group; E, n = 7-8) or t-test (B & D, n = 10-17 mice/ group). Data are shown as mean ± SEM.

### FGF21 was required for the increased consumption of high-protein dessert following a LP pre-load meal

To determine the contribution of FGF21 to these outcomes, we repeated the experiment using male (Fig. 4A-C) and female (Fig. 4D-E) FGF21^ΔL^ mice and control littermates. Consistent with the understanding that plasma FGF21 derives primarily from the liver, LP-feeding increased plasma FGF21 levels in littermates, but not in FGF21^ΔL^ mice [p (genotype x diet) < 0.001 (Fig. 4C)]. As before, protein-restricted mice and controls consumed most of their total caloric intake from maintenance diet, during the 2-h preload meal [p (time)< 0.001, Tukey’s posthoc, p< 0.001 (Fig. 4A, D)]. This did not depend on genotype (Fig. 4A, D). In agreement with the previous findings, protein-restricted male mice ate more of the high-protein dessert compared to control littermates [p (pre-load diet treatment) < 0.05, Tukey < 0.01]. This was significantly blunted in mice lacking liver FGF21, compared to littermate controls [p (pre-load diet treatment) < 0.05; Tukey, p< 0.05 (Fig.4B)]. Consistent with previous results, there was no effect of either protein restriction or genotype on consumption of high-protein dessert in female mice (Fig. 4E).

**Figure 4.**
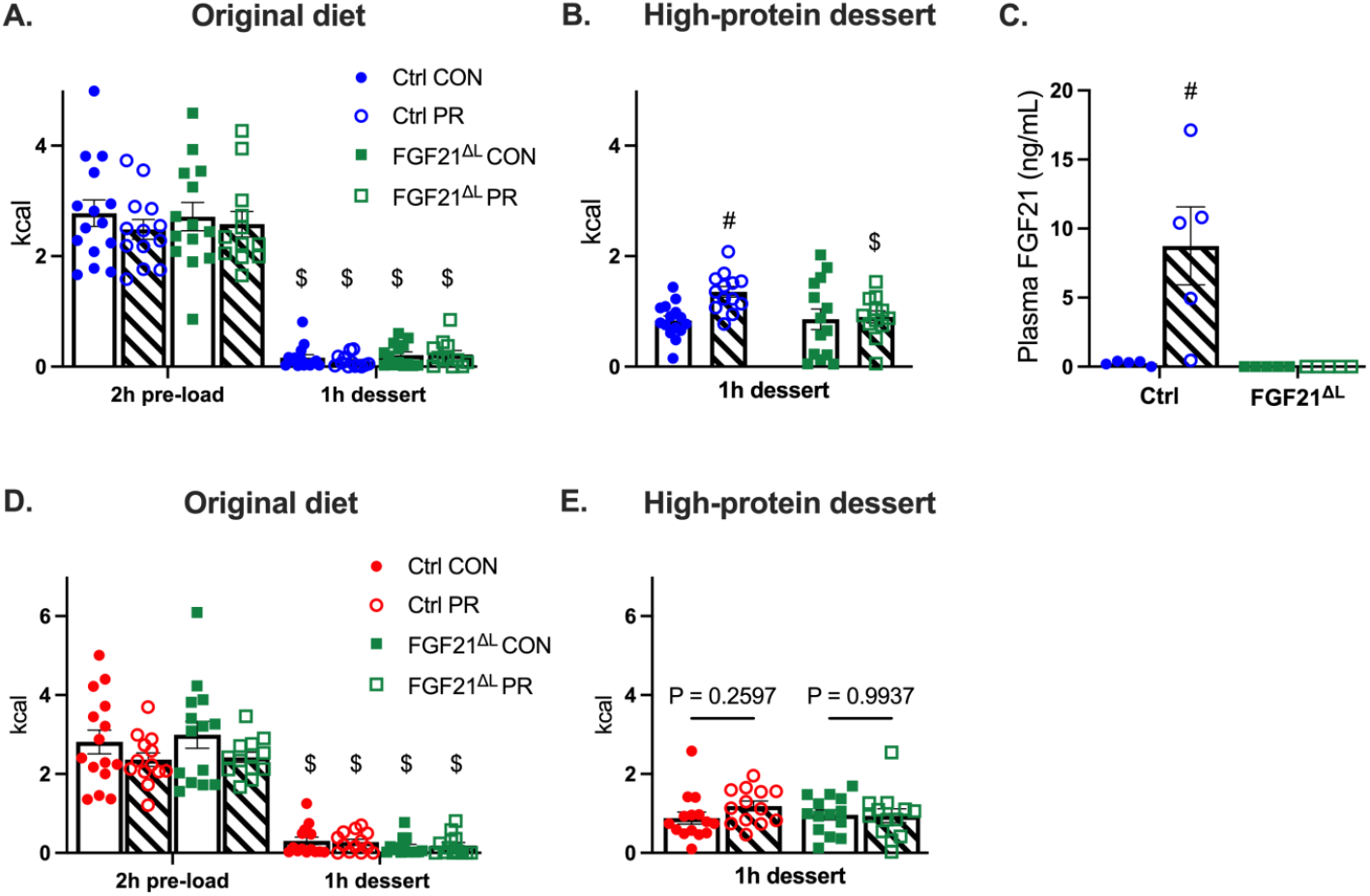
Liver FGF21 is required for the increased consumption of high-protein dessert in protein-restricted male mice. Both male (A-C) and female (D,E) CON and PR mice consumed most of their caloric intake from original maintenance diets during the 2h pre-load meal and ate relatively little pelleted diet during the ‘dessert’ phase of the experiment [$: p<0.0001 versus 2h pre-load] (A, D). After the pre-load, PR male littermates ate significantly more high-protein dessert compared to CON male littermates, and this was significantly blunted in FGF21^ΔL^ mice [#: p<0.01 versus Ctrl CON; $: p<0.05 versus Ctrl PR] (B). Plasma FGF21 was robustly increased in PR male littermates but not in PR male FGF21^ΔL^ mice (C). There was no effect of protein restriction on high-protein ‘dessert’ intake in female mice (E). Analyses were made by two-way ANOVA with Tukey *post hoc* tests (A-B, n = 14-15 mice/ group; C, n = 5 mice/ group; D-E, n = 13-15 mice /group) Data are shown as mean ± SEM.

## Discussion

A higher intake of sugar-sweetened beverages and desserts has been consistently associated with overweight and obesity [1,2] and related metabolic disorders [4–7]. However, adherence to the recommendations to avoid added sugars and processed sweet treats remains poor [8,9], resulting in an ongoing increase in sugar consumption [10]. Recent studies demonstrated that FGF21 is secreted from the liver in response to dietary protein restriction [11–15,23]. Furthermore, pharmacologic administration of FGF21 acts via the nervous system to decrease sugar intake [18,19]. Together these data suggested a nutritional strategy to reduce the motivation for eating sweet desserts, by reducing protein content of the preceding meal. Accordingly, we hypothesized that eating a low-protein pre-load meal would decrease the subsequent consumption of unhealthy sweet desserts. Contrary to expectation, however, we observed no effect of dietary protein restriction on the consumption of either liquid or solid sucrose dessert following a pre-load meal.

A history of dietary protein restriction did not influence the consumption of sucrose dessert in calorically sated mice. To account for potential differences in palatability, or in metabolic outcomes that depend on concentration or substrate form [35,36], we repeated the study four times. We varied the concentration of sucrose solution; this had no effect on the outcomes. We used solid sucrose pellets; this also had no effect on the outcomes. These results surprised us for several reasons. First, we and others previously observed a robust effect of pharmacologic FGF21 treatment, or its transgenic overexpression, to reduce sucrose intake in *ad libitum* fed individuals [17–19,24,32,37]. Second, our previous studies demonstrated that as few as 4 days of low protein feeding robustly increases endogenous plasma FGF21 in mice [12]. Lastly, 3 days of dietary protein restriction robustly increased plasma FGF21 in this experiment—yet it was not sufficient to alter sucrose intake. Importantly, however, the rise in plasma FGF21 observed following pharmacologic or transgenic approaches was considerably higher than we have observed in response to protein restriction. For example, transgenic mice overexpressing FGF21 have a plasma concentration of roughly 500 ng/mL [19]. When Larson and colleagues injected 0.1mg/kg hFGF21 to male mice, we observed a peak plasma concentration of roughly 65 ng/mL [17]. By contrast, the 3 days LP diet treatment in this study resulted in only about 5-10 ng/mL plasma FGF21. A likely interpretation of our findings, therefore, is that dietary protein restriction does not induce sufficient endogenous FGF21 to alter sucrose intake in sated mice.

FGF21 secretion in response to protein restriction in this study was sufficient to influence the subsequent intake of a protein-rich dessert or snack. Male mice consuming a low P: E pre-load meal exhibited a significant increase in the consumption of a high-protein ‘dessert’ or snack; this effect was attenuated in male mice lacking hepatic FGF21. These findings align with previous evidence demonstrating that human subjects eating a low-protein breakfast acquired a conditioned flavor preference for a subsequent high-protein (but not low-protein) lunch, and this was abrogated with a high-protein breakfast [38]. Likewise, prior studies report that a history of protein-restriction induced a stronger preference for high-protein solutions in rodents [11,39,40], and that endogenous FGF21 induced by protein restriction facilitates subsequent liquid protein intake in rodents [11]. Our findings extend this understanding to include solid protein sources (rather than 4% casein solution) offered to calorically sated mice (rather than fasted or *ad libitum fed*). They are the first to identify liver as the required endogenous source of FGF21 modulating macronutrient preference, since previous studies used whole-body FGF21 knockout mice [11]. Continued food intake in calorically sated individuals is generally interpreted as reflecting reward-driven or hedonic feeding [27–30]. Thus, additional research focused on how liver-derived FGF21 alters dopaminergic neurotransmission [24,40,41] and state-dependent motivated behaviors like place preference and operant responding are warranted [25,26,42].

In agreement with previous reports of sex differences in macronutrient selection [43–45] and FGF21 biology [12,23,46], we observed a differential response between male and female mice in this study. Specifically, we did not observe an effect of protein restriction on subsequent consumption of a high protein ‘dessert’ in female mice. Although both male and female mice increased plasma FGF21 after 3 days of protein restriction, the effect was significantly more robust in males. And although we previously reported that female mice treated with pharmacologic FGF21 shift macronutrient selection towards increased consumption of dietary protein, the magnitude of this outcome is significantly blunted compared to male counterparts [37]. Therefore, the sex-dependent behavior observed in the present study likely depends on sex differences in both FGF21 secretion and FGF21 sensitivity. Regarding sex differences, the current findings contrast with a recent report showing a history of protein restriction increases the preference for casein vs maltodextrin solutions a 2-bottle choice test in both male and female mice [39]. Thus, female mice *can* adjust feeding behavior and macronutrient selection in response to protein status, but this like depends on specifics of the experimental design. Some important differences between the two studies include feeding status (calorically sated vs *ad libitum fed*), protein source (Kool-Aid flavored solution vs. unflavored, solid diet), and the length of protein restriction leading into the test (3 vs ∼18 days). Lastly, our experimental design did not account for estrous cycle—yet cycle stage did not affect protein preference in the previous study [39].

## Conclusion

Our study sheds additional light on the relationship between FGF21 and eating behavior. Contrary to our expectation, although a history of dietary protein restriction was sufficient to dramatically increase circulating FGF21 in male and female mice, this did not reduce the subsequent consumption of sweet desserts. Thus, pharmacologic doses of this hormone may be necessary to modulate the motivation to consume sweet foods in calorically sated individuals and nutritional strategies to induce FGF21 may be ineffective. By contrast, a history of protein restriction did significantly increase the consumption of a protein-rich snack in otherwise calorically sated male mice, and this depended on secretion of FGF21 from the liver. These findings are consistent with a growing literature that link FGF21 as a critical mechanism underlying reward-based protein intake. Understanding the underlying mechanisms could have significant implications for developing behavioral strategies to harness FGF21 signaling to promote healthy eating habits.

## Acknowledgements

This work was supported by the National Institutes of Health R01DK121035 to KKR. CTW was supported by the Yen Chuang Taiwan Fellowship from the University of California, Davis. KRL was supported by a University of California, Davis Floyd and Mary Schwall Dissertation Fellowship. LCS was supported by the National Institute of General Medical Sciences funded training program NIH T32GM099608. We thank Nadejda Godoroja and Yanbin Fang for excellent technical assistance.

---

**Supplementary Figure 1.**
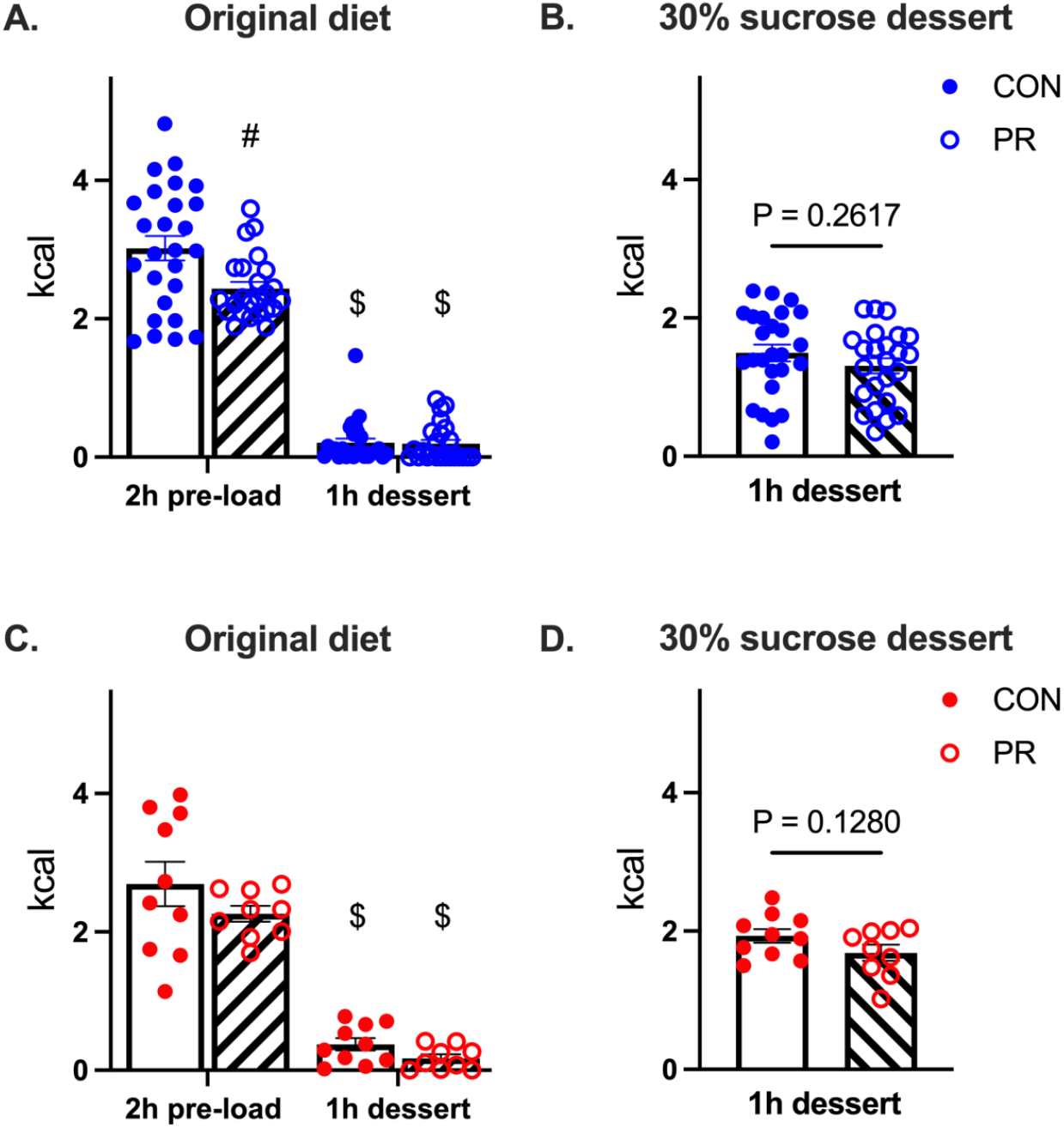
Protein status of a pre-load meal did not affect subsequent consumption of 30% sucrose dessert. Male (A-B) and female (C-D) CON and PR mice consumed the majority of their caloric intake from original maintenance diets during the 2h pre-load meal and ate relatively little pelleted diet during the ‘dessert’ phase of the experiment [$: p<0.0001 versus 2h pre-load; #: p<0.001 versus CON] (A, C). There was no effect of pre-load diet on the consumption of 30% sucrose dessert (B, D). Analyses made by two-way ANOVA with Tukey *post hoc* tests (A,C) or by T-test (B,D). Data are shown as mean ± SEM, n= 9-18 mice per group.

**Supplementary Figure 2.**
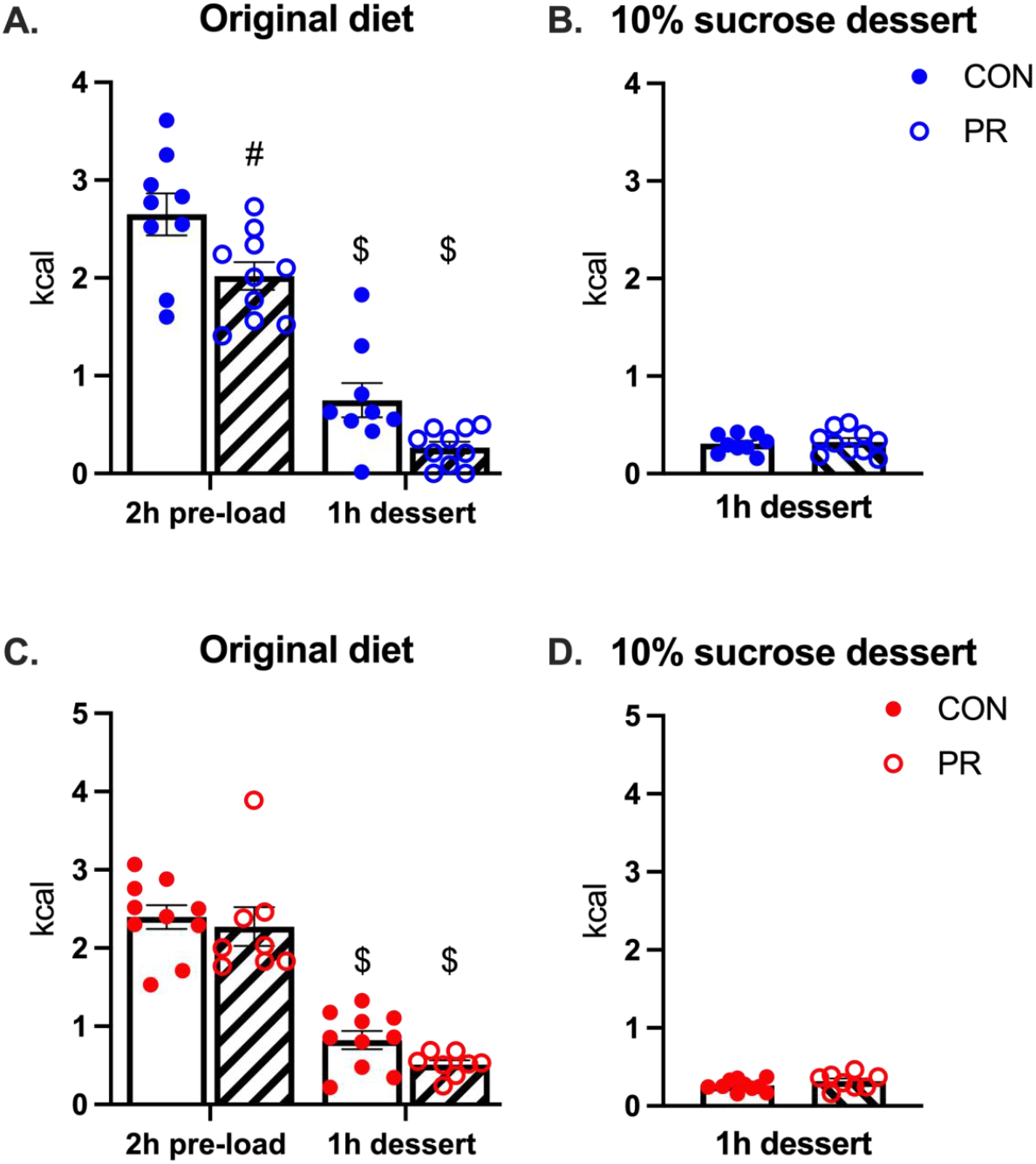
Protein status of a pre-load meal did not affect subsequent consumption of 10% sucrose dessert. Male (A-B) and female (C-D) CON and PR mice consumed the majority of their caloric intake from original maintenance diets during the 2h pre-load meal and ate relatively little pelleted diet during the ‘dessert’ phase of the experiment [$: p<0.0001 versus 2h pre-load; #: p<0.001 versus CON] (A, C). There was no effect of pre-load diet on the consumption of 10% sucrose dessert (B, D). Analyses made by two-way ANOVA with Tukey *post hoc* tests (A,C) or by T-test (B,D). Data are shown as mean ± SEM, n= 8-10 mice per group.

**Supplementary Figure 3.**
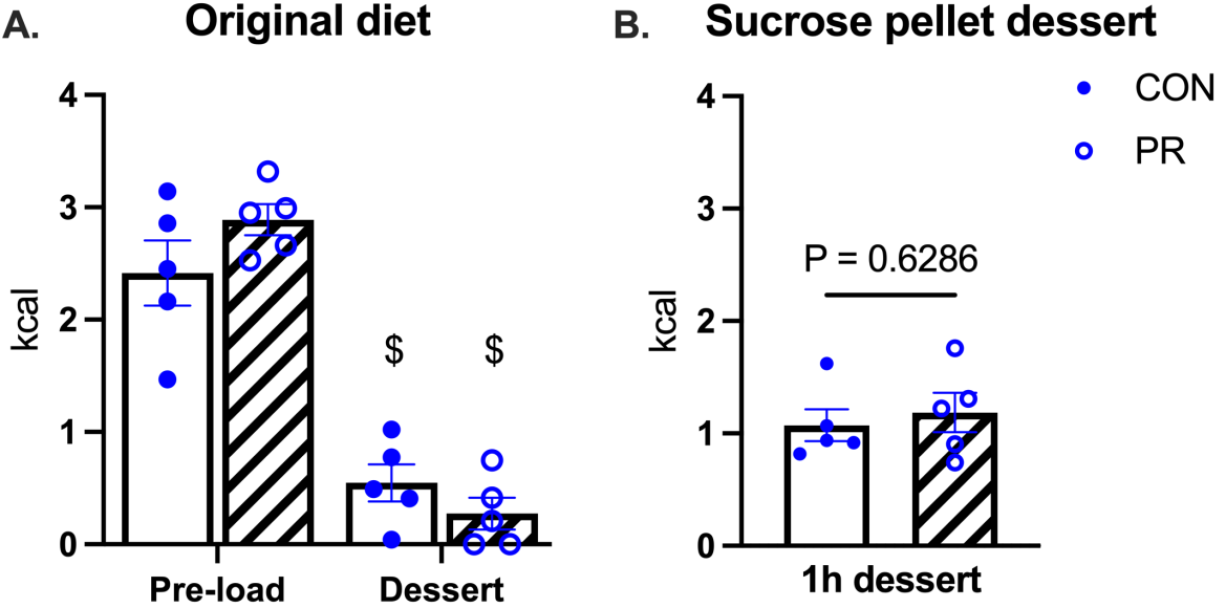
Protein status of a pre-load meal did not affect subsequent consumption of sucrose pellets. CON and PR male mice consumed the majority of their caloric intake from original maintenance diets during the 2h pre-load meal and ate relatively little pelleted diet during the ‘dessert’ phase of the experiment [$: p<0.0001 versus 2h pre-load] (A). However, there is no effect of pre-load diet on the consumption of sucrose dessert (B). Analyses made by two-way ANOVA with Tukey *post hoc* tests (A,C) or by T-test (D,E). Data are shown as mean ± SEM, n= 5 mice per group

